# Household geckos as a potential vector for *Salmonella* transmission in Malawi

**DOI:** 10.1101/2023.09.08.556805

**Authors:** Catherine N. Wilson, Patrick Musicha, Mathew A. Beale, Yohane Diness, Oscar Kanjerwa, Chifundo Salifu, Zefaniah Katuah, Patricia Duncan, John Nyangu, Andrew Mungu, Muonaouza Deleza, Lawrence Banda, Nicola Elviss, Christopher P. Jewell, Gina Pinchbeck, Nicholas R. Thomson, Nicholas A. Feasey, Eric M. Fèvre

## Abstract

*Salmonella* was isolated from 23/79 (29.1%) gecko stool samples from households in southern Malawi. Whole genome sequencing of 47 isolates revealed 27 *Salmonella* serovars spanning two subspecies. Our results demonstrate that geckos play an important role in the carriage and potentially, transmission of *Salmonella* within households.

## Introduction

*Salmonella* is a major cause of foodborne disease globally and this zoonosis represents a concern to both human and animal health. *Salmonella* can be carried asymptomatically within the stool of humans, in animals and within the environment. *Salmonella enterica* subspecies *enterica* is consistently associated with disease in warm-bloodied animals, while non-enterica subspecies of *Salmonella enterica* have been associated only with opportunistic infection in humans (1,2). Some of the *S. enterica* serovars, including *S*. Typhimurium and *S*. Enteritidis, have a broad host range. Others are associated with disease in a single species but are also, less commonly, associated with disease in other animal species. Host-restricted serovars only cause disease within a single species. This means that ecologically, characterisation of the *Salmonella* spp. present within a household is important to gauge the potential zoonotic risk for other species present.

The African house gecko (*Hemidactylus mabouia*) is a species of gecko widely distributed across the neotropics and sub-Saharan Africa (3). The species is primarily nocturnal, although some geckos may explore during the day. The diet of geckos is a varied insectivorous diet including isopods, centipedes, spiders, scorpions, cockroaches, beetles, moths, flies, mosquitoes, anoles and other geckos (4).

Free-living and captive reptiles have been routinely identified as reservoirs of *Salmonella* spp. (5–7). Reptiles have been previously found to carry a number of serovars, some of which are linked to invasive disease in humans (8). Previously, a high prevalence (23.9%) of *Salmonella* carried within gecko faeces has been described in Asia (9). This is the first time in 30 years that carriage of *Salmonella* by geckos free-living within a household environment in Africa has been investigated (10).

### The Study

Thirty households were sampled at three time points over the period of a year (November 2018-December 2019) in both an urban (Ndirande) and a rural (Chikwawa) study site in southern Malawi to investigate carriage of *Salmonella* by humans, by animals and within the environment. Assistant Veterinary Officers, who carried out the sampling, were appropriately trained by the Study Principal Investigator to recognise gecko faeces. Characteristic samples of gecko faeces (Figure 1) were collected from the walls of households within the study. If multiple gecko stool were collected from a single household at a single visit these samples were pooled prior to further processing.

**Figure 1:**
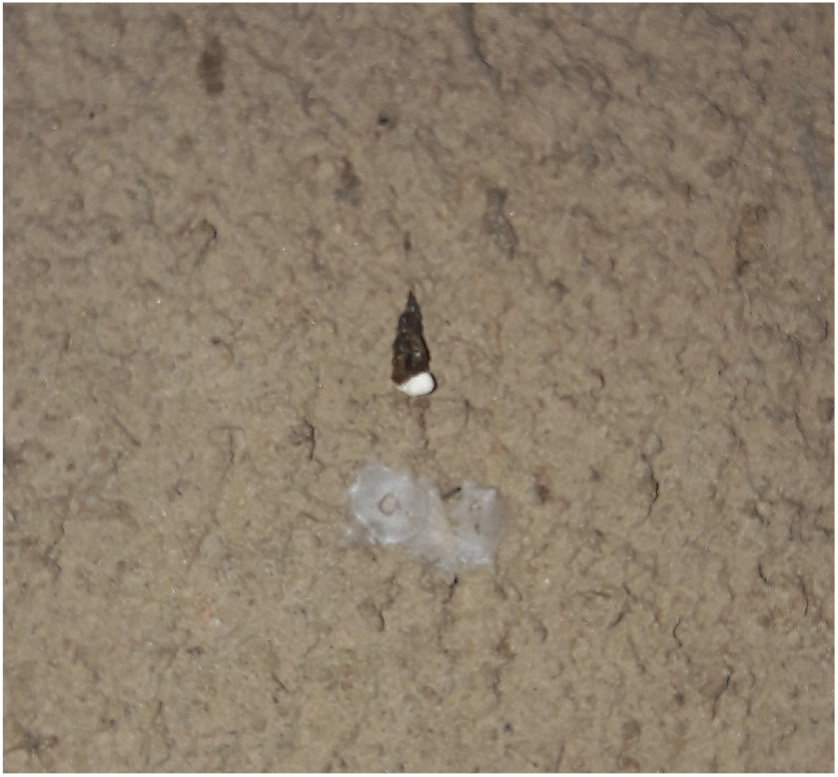
Characteristic gecko faeces; stool has a cylindrical, tapered shape, roughly 1.5cm long, mainly brown coloured stool with a small white portion which contains urate excretions.

To detect *Salmonella*, selective microbial culture was undertaken using XLD and Harlequin chromogenic agar for *Salmonella* esterase (CASE). Anti-sera testing against the O antigen and qPCR analysis using the *ttr* primer were used to detect and confirm the presence of *Salmonella* and a subset of the isolates (two picks from each sample) were submitted for whole genome sequencing (11).

Ethical approval for this study was obtained from the College of Medicine Ethics Committee (Malawi) reference number P.02/18/2368) and the University of Liverpool Research Ethics Committee (Reference number VREC686). Informed consent from household owners was obtained for the collection of gecko faeces from each household at each visit, and for the other samples collected.

Salmonellae were isolated from 23/79 (29.1%) of the gecko stool samples. *Salmonella* positivity rate was by far higher in gecko samples from Chikwawa (21/47 (44.7%)), the rural study site, than Ndirande, the urban study site (2/32 (6.3%)).

Between two to three colony picks of *Salmonella* from each sample were submitted for whole genome sequencing. Forty-seven good quality *Salmonella* genomes were obtained from geckos including forty-two from Chikwawa and five from Ndirande; representing whole genome sequences from 20 discrete gecko stool samples. Of these 20 gecko samples, 19 were from Chikwawa and 1 was from Ndirande. Within sample diversity was detected in seven samples; where two different *Salmonella* serovars were detected from a single sample following whole genome sequencing.

There were twenty-seven unique genomes in the collection of 47 genomes covering 2 subspecies of *Salmonella; S. enterica* and *S. salamae* (Table 1). The genome isolated from Ndirande was *S. enterica* subspp. *Oranienburg*. The genomes isolated from Chikwawa were 11 *S. enterica* covering nine serovars, and 15 *S. salamae* covering 13 different antigenic profiles. Of these 27 different serovars, disease in humans has previously been reported to be caused by ten of these serovars (37.0%).

**Table 1:** Results of whole genome sequencing of *Salmonella* isolates collected from geckos, including genotypic and phenotypic antimicrobial resistance profiles. CHH*/NHH* = individual household number prefix, ukn = unknown, AMR = antimicrobial resistance, S = sensitive, I = intermediate, R = resistant, NA = not applicable.

| Study site | Household number | Visit number | Sample ID | Subspecies | Serovar | ST | Genotypic AMR | Phenotypic AMR |  |  |  |  |  |
| --- | --- | --- | --- | --- | --- | --- | --- | --- | --- | --- | --- | --- | --- |
|  |  |  |  |  |  |  |  | Tetracycline | Nalidixic acid | Chloramphenicol | Cefpodoxime | Streptomycin | Ampicillin |
| Chikwawa | CHH1 | 1 | 21 | salamae | II 1,13,23:z29:e,n,x | 1188 | none | S | S | S | S | I | R |
| Chikwawa | CHH1 | 2 | 20 | salamae | II 1,4,12,27:-:1,5 | ukn | *fosA7* | S | S | S | S | I | S |
| Chikwawa | CHH1 | 1 | 21 | salamae | II 35:z29:1,5,7 | ukn | *fosA7* | S | S | S | S | S | S |
| Chikwawa | CHH1 | 2 | 20 | enterica | Concord | 694 | none | S | S | S | S | S | S |
| Chikwawa | CHH2 | 2 | 5 | salamae | II 1,4,12,27:-:1,5 | ukn | *fosA7* | S | S | S | S | I | S |
| Chikwawa | CHH2 | 3 | 21 | enterica | Anatum | 64 | none | NA | NA | NA | NA | NA | NA |
| Chikwawa | CHH2 | 3 | 21 | enterica | Gaminara | 2152 | none | NA | NA | NA | NA | NA | NA |
| Chikwawa | CHH2 | 3 | 5 | enterica | Kingabwa | 546 | *fosA7.7* | S | S | S | S | I | S |
| Chikwawa | CHH3 | 3 | 13 | enterica | Jangwani | 3918 | none | NA | NA | NA | NA | NA | NA |
| Chikwawa | CHH4 | 2 | 20 | salamae | II 1,13,23:z29:- | ukn | none | NA | NA | NA | NA | NA | NA |
| Chikwawa | CHH4 | 3 | 18 | enterica | Poona | 2566 | none | S | S | S | S | S | S |
| Chikwawa | CHH5 | 3 | 6 | salamae | II 41:I,[z13],[z28]:z39 | ukn | none | S | S | S | S | I | S |
| Chikwawa | CHH5 | 1 | 22 | enterica | Ahuza | 2215 | *fosA7.7* | R | S | S | S | S | S |
| Chikwawa | CHH6 | 1 | 20 | salamae | II 35:g,z62:- | ukn | none | S | S | S | S | S | S |
| Chikwawa | CHH7 | 2 | 13 | salamae | II 39:a:1,5,7 | ukn | none | S | S | S | S | S | S |
| Chikwawa | CHH8 | 1 | 18 | salamae | II 1,4,12,27:e,n,x:e,n,x | ukn | *fosA7* | S | S | S | S | I | S |
| Chikwawa | CHH8 | 1 | 18 | enterica | Gaminara | 2152 | none | R | S | S | S | I | S |
| Chikwawa | CHH9 | 2 | 22 | enterica | Enteritidis | 366 | none | R | S | S | S | S | S |
| Chikwawa | CHH11 | 3 | 14 | enterica | Bukavu | 5410 | none | S | S | S | S | S | S |
| Chikwawa | CHH12 | 3 | 18 | salamae | II 6,7:z:z6 | ukn | none | S | S | S | S | R | S |
| Chikwawa | CHH12 | 3 | 18 | enterica | Gaminara | 2152 | none | S | S | S | S | S | S |
| Chikwawa | CHH14 | 2 | 16 | salamae | II 1,9,12,46,27:b:1,5 | ukn | none | S | S | S | S | S | S |
| Chikwawa | CHH15 | 1 | 8 | salamae | II 6,7:z:z6 | ukn | none | S | S | S | S | I | S |
| Chikwawa | CHH15 | 3 | 6 | salamae | II 35:g,z62:e,n,x | ukn | *fosA7* | S | S | S | S | S | S |
| Chikwawa | CHH15 | 2 | 19 | salamae | II 1,13,23:e,n,x:e,n,x | ukn | none | S | S | S | S | S | S |
| Chikwawa | CHH15 | 2 | 19 | salamae | II 1,13,23:-:e,n,x | ukn | none | S | S | I | S | I | S |
| Ndirande | NHH6 | 3 | 9 | enterica | Oranienburg | ukn | *fosA7* | S | S | S | S | S | S |

Genotypic antimicrobial resistance determinants were carried by 8/27 (29.6%, 95% CI 15.8-48.5%) of the unique genomes, seven from Chikwawa and 1 from Ndirande. Six of the eight AMR determinants were *fosA7*, two were a variant of the *fosA7* gene; *fosA7*.*7*. These AMR determinants were all chromosomally integrated.

## Discussion and conclusions

Here we present for the first time a characterisation of *Salmonella* carried by household geckos in Malawi, demonstrating high carriage rates of *Salmonella* within gecko stool. Some of the serovars carried by geckos may have the potential for pathogenicity to both animals and humans, eg *S*. Enteritidis, *S*. Poona and *S*. Concord.

We have found a significantly higher carriage of *Salmonella* within rural households in comparison to those located in an urban area. The reasons for this are unclear and may be related to climatic factors; the rural area has a higher mean average temperature (24.2°C) than the urban study site (20.6°C)(12). It may also be related food availability for the geckos; flies and insects. In the rural area there are a greater number of livestock and more vegetation, which could lead to a greater profusion of the insect population. However, further work is needed to investigate this in more depth, and explore the ecological niches exploited by geckos that may lead to *Salmonella* exposure.

Some of the *Salmonella* serovars detected have the potential to cause invasive and non-invasive disease in humans and/or animals, as documented in other published literature (13– 18). In this study, humans and animals were sampled simultaneously and there were no reports of clinical disease affecting any of the sampled hosts, therefore it was presumed all *Salmonella* carriage was asymptomatic.

Geckos play an important role in household ecology, controlling insect populations with little disturbance to human and animal inhabitants of households. At the same time, they freely move around the household environment, leaving stool on surfaces which humans may come into contact with. The finding that geckos carry serovars which have the potential to cause pathogenic disease in other species in sub-Saharan Africa has important implications for public health and adds to the known picture of movement of *Salmonella* bacteria around the household, providing a deeper understanding of the ecology of this bacterial species.

This is not a call to arms against the gecko population and any population control targeting the species at a household level is certainly not the recommendation made here. Rather, we should consider that geckos may act as a sentinel of *Salmonella* within households. It is likely that geckos are colonised by *Salmonella* following consumption of contaminated insects or environmental material around the household and so the prevalence of *Salmonella* detected within gecko stool may reflect the overall level of *Salmonella* contamination of the household environment.

It is important that hygiene precautions are taken within households where geckos are present to minimise the likelihood of transmission of *Salmonella* bacteria to other animals or humans and their environment. Important interventions to achieve this and so decrease the risk of faeco-oral bacterial transmission include good hand hygiene (particularly prior to preparation of food or following use of the bathroom), regular sweeping and cleaning of living spaces, keeping rubbish off the floor, keeping dishes and utensils clean and ideally in a sealed container between usages and cleaning the household with appropriate and effective disinfectants. These actions should help to reduce the risk of zoonotic transmission of *Salmonella* between geckos and humans.

## Acknowledgement

We would like to thank the households recruited to this study for their participation in the sample collection. This work has been supported by a Wellcome Trust Clinical Fellowship award to Catherine N. Wilson, grant number 203919/Z/16/Z.

## Notes

### Competing Interest Statement

The authors have declared no competing interest.

